# High heritability of telomere length, but low evolvability, and no significant heritability of telomere shortening in wild jackdaws

**DOI:** 10.1101/2020.12.16.423128

**Authors:** Christina Bauch, Jelle J. Boonekamp, Peter Korsten, Ellis Mulder, Simon Verhulst

## Abstract

Telomere length (TL) and shortening rate predict survival in many organisms. Evolutionary dynamics of TL in response to survival selection depend on the presence of genetic variation that selection can act upon. However, the amount of standing genetic variation is poorly known for both TL and TL shortening rate, and has not been studied for both traits in combination in a wild vertebrate. We used experimental (cross-fostering) and statistical (animal models) means to disentangle and estimate genetic and environmental contributions to TL variation in pedigreed free-living jackdaws (*Corvus monedula*). TL was measured twice early in life (age- and interval-standardized), when shortening is highest, using the high-precision TRF technique, adapted to exclude interstitial telomeric sequences. TL shortened significantly during the nestling phase (10.4 bp/day), was highly repeatable within individuals (*R*=0.97) and genetically correlated between the two ages (*r*_G_>0.99). Additive genetic effects explained the major part of TL variation between individuals, with heritability on average estimated at *h*^*2*^=0.74. Parent-offspring regressions yielded similar estimates for the sexes when accounting for changes in paternal TL over life. Cohort effects explained a small but significant part of TL variation. Heritable variation for telomere shortening was negligible. Despite the high heritability of TL, its evolvability, which scales the additive genetic variance by mean TL, was close to zero. Hence evolutionary change of TL is likely to be slow despite significant selection.

## Introduction

Individuals differ in lifespan and other fitness components, and identification of the molecular and physiological traits associated with fitness is a useful step in deciphering ecological and evolutionary dynamics of life histories. Telomere length (TL) is such a trait, because shorter TL predicts lower survival in humans (Boonekamp, Simons, Hemerik, & Verhulst, 2013) and wild vertebrates (Wilbourn et al., 2018). TL is therefore increasingly used as biomarker of ageing and phenotypic quality across research areas such as epidemiology, ecology and evolutionary biology (Monaghan et al. 2018). Telomeres consist of evolutionarily conserved, non-coding DNA sequence repeats (Blackburn, 1991) that together with the shelterin protein complex form the ends of chromosomes (de Lange, 2005) and contribute to genome stability (O’Sullivan & Karlseder, 2010). Telomeres are dynamic structures, in that their length shortens with age due to incomplete replication during cell division, which can be accelerated by DNA- and protein-damaging factors and attenuated or counteracted by maintenance processes (Chan & Blackburn, 2004).

To understand how TL may play a role in shaping life-histories from an evolutionary perspective, it is necessary to quantify its additive genetic variance and resulting heritability that natural selection can act upon. However, heritability estimates of TL differ largely between studies, and to what extent variation in TL is due to inheritance or whether it is mainly driven by the environment, is therefore still under debate (Atema et al., 2015; Benetos et al., 2019; Broer et al., 2013; Dugdale & Richardson, 2018). Differences in the genetic background of populations, temporal-spatial variation of environmental influences, and methodological differences including limitations to separate genetic and environmental effects can all underlie variation in heritability estimates between studies (Becker et al., 2015, Dugdale & Richardson, 2018).

TL at any given age is defined by the initial TL as transferred through the parental gametes and the TL change from the zygote stage onwards, and, like TL itself, TL dynamics varies under the influence of both genes and the environment. For example, TL generally shortens with increased exposure to various (environmental) stressors (Angelier, Costantini, Blévin, & Chastel, 2018; Bauch, Becker, & Verhulst, 2013; Boonekamp, Mulder, Salomons, Dijkstra, & Verhulst, 2014; Hall et al., 2004; Kotrschal, Ilmonen, & Penn, 2007; McLennan et al., 2016; Young, Barger, Dorresteijn, Haussmann, & Kitaysky, 2013), and telomere shortening is typically faster early in life (Benetos et al., 2019; Salomons et al.; 2009, Spurgin et al., 2018). Thus, exposure to environmental influences, age-dependent sensitivity to environmental effects and age-dependent gene expression all contribute to TL variability. Most studies estimating TL heritability, however, measured TL at different ages in different individuals (reviewed in Dugdale & Richardson, 2018). Not taking this increased TL variability into account, or doing so only by statistically controlling for age effects will likely lead to underestimated TL heritability. On the other hand, a shared environment among related individuals can increase their phenotypic similarity and thereby inflate heritability estimates (Kruuk & Hadfield, 2007). Parental effects can potentially influence TL in multiple ways, via additive genetic effects on gamete TL and TL regulation, epigenetic effects (e.g. mediated by paternal age; Bauch, Boonekamp, Korsten, Mulder, & Verhulst, 2019; Eisenberg, 2019; Noguera, Metcalfe, & Monaghan, 2018), and via early-life parental effects of genetic and non-genetic origin. Measuring TL inheritance is therefore challenging.

‘Animal models’ are a powerful statistical tool to disentangle genetic from environmental effects, which makes use of all relatedness information available in a pedigree (Wilson et al., 2010). Several studies investigated TL heritability in wild vertebrates using animal models (e.g. Asghar, Bensch, Tarka, Hansson, & Hasselquist, 2015; Becker et al., 2015; Sparks et al., 2020; Voillemot et al., 2012), but these studies all measured TL using qPCR (Nussey et al., 2014). This technique pools telomeres at the end of chromosomes with the interstitial telomeric sequences (ITS) in the TL estimates, and the latter are frequent in many genomes of species other than humans, and can vary strongly in number and length between individuals (Foote, Vleck, & Vleck, 2013; Meyne et al., 1990). It is not clear therefore how the currently available estimates of the heritability of pooled estimates of all telomeric sequences in the genome relate to the heritability of the length of terminal telomeres. Even less is known of genetic variation in telomere shortening, with one study on human twins reporting a heritability of 0.28 (Hjelmborg et al., 2015).

Heritability (*h*^2^) is estimated as the ratio between additive genetic variance (V_A_) and the total phenotypic variance (V_P_) *h*^2^=V_A_/V_P_ (Falconer & Mackay, 1996) and as such, low heritability can result from low additive genetic variance, but also from high total variance, e.g. through environmental effects. The evolutionary potential of a trait is dependent on the additive genetic variance, which scaled to the mean of the trait yields a metric known as the evolvability (Houle, 1992). We are aware of one study estimating the evolvability of TL, in an insect, which found evolvability to be low despite high heritability (Boonekamp et al., 2020a).

We estimated the additive genetic and environmental contributions to the variability of TL among and within individuals to estimate TL heritability, the age-genotype interaction and the evolutionary potential in a population of jackdaws *Corvus monedula* with a known multigenerational pedigree. We measured TL in erythrocytes, using the golden-standard technique telomere restriction fragment analysis (TRF) adapted to exclude ITS (Salomons et al., 2009). Individuals were sampled twice early in life, when TL shortening is highest, at the same age and with a fixed time interval between the first and second sampling. To disentangle genetic and environmental causes of variation in TL and TL shortening we applied (a) experimental cross-fostering and (b) animal model analysis (Wilson et al., 2010), and used the estimates to calculate the evolvability of TL. We additionally performed parent-offspring regressions, and assessed the relative maternal and paternal contributions to TL inheritance, taking also effects of paternal age at conception into account (Bauch et al., 2019).

## Material and methods

### Study population

We studied a jackdaw *Corvus monedula* population breeding in nest box colonies located south of Groningen, The Netherlands (53°14’N, 6°64’E), which has been under investigation since 1996. Birds in our study population are ringed with numbered metal rings and colour rings shortly before fledging or as immigrants at first breeding in the study nest boxes. Breeders are highly site faithful and all breeding birds are identified by their unique colour ring combinations via telescope, photo or video camera. Jackdaws breed monogamously with low divorce rates and very rare extra-pair paternity (Henderson, Hart, & Burke, 2000; Liebers & Peter, 1998). However, due to partner death, about 50% of the adults in our dataset had two or more partners and the population comprises full as well as half-siblings produced over multiple years. From one day before the expected hatching date the nest boxes were checked daily for hatchlings. Freshly hatched chicks were marked by specific combinations of clippings of the tips of the toenails for identification until ringing. Exact ages were known for jackdaws native to the study’s nest boxes. Immigrants were assigned an age of 2 years when breeding for the first time, which is the modal age at recruitment in our population. The sex of jackdaws has been identified either molecularly (Griffiths, Double, Orr, & Dawson, 1998) or by behavioural observations and cross-reference with breeding partners.

### Pedigree

The pruned pedigree consists of 1007 relatedness-informative individuals, whereof 715 individuals held data on TL at the age of 4 days and 474 individuals additionally TL data when 29 days old. The pedigree spans 6 generations for known TL of either age. For details on relationships in the pedigrees see table S1, fig. S1.

### Cross-fostering

To disentangle additive genetic from early-life parental effects we performed cross-fostering manipulations in our study population. In 2015 and 2016 in a subset of the nest boxes (n=58) complete clutches were exchanged between nests during mid incubation. Nests were matched for clutch size and laying date (±1 day), but in all other respects cross-fosterings were performed randomly. In all years, brood sizes were manipulated in most nests and nestlings were transferred between nests when the oldest nestling was 4 days old. Age-matched broods were reduced by removing 3 chicks and adding one chick and enlarged by adding 3 chicks and removing one. Consequently, manipulated broods (i.e. between first and second TL measurement) comprised genetic and non-genetic offspring of parents and thus, both genetic and non-genetic siblings. For details, see (Boonekamp, Bauch, & Verhulst 2020; Boonekamp et al., 2014). The multigenerational pedigree included individuals with various degrees of relatedness which experienced the same or different rearing environments. The cross-fostering experiments further lead to additional information in the dataset by including full siblings that experienced different environments, e.g. number and identity of siblings or parental care. The proportion of cross-fostered offspring by clutch exchange was 9% and by exchange at the age of 4 days was 33%.

### Blood sampling and telomere analysis

TL was measured in blood samples taken between 2005-2016 from 715 individuals at the maximum age of 4 days (mean±SD=3.7±0.6 days) and again 25 days later in a subset of 474 individuals (age 29 days). This interval between repeated telomere measurements of nestlings covers approximately 70% of their time in the nest (Röell, 1978). Fathers were caught and blood sampled later in the breeding seasons when the brood was at least 14 days old, contributing telomere data of 82 fathers at conception across multiple years to the dataset. Samples were stored in 2% EDTA buffer at 4-7 °C and within 3 weeks snap frozen in a 40% glycerol buffer for permanent storage at −80 °C. TL of nucleated erythrocytes was measured performing TRF analysis under non-denaturing conditions as in Salomons et al. (2009). See supplementary material for details. With this method, only terminally located telomere sequences were quantified and ITS were excluded. A sample of an individual consists of a characteristic TL distribution from erythrocytes of different ages and different chromosomes within cells. The individual average of this TL distribution was used for further analyses (Salomons et al., 2009). Samples were run on 57 gels. Samples from chicks of the same brood were partly spread over different gels but repeated samples from individuals were run on the same gel. The coefficient of variation of one control sample from a 29 days old jackdaw run on 26 gels was 6% and of one control sample of a goose, with a similar TL distribution in a similar range, run on 31 other gels was 7%.

### Statistical analyses

First, we investigated some descriptive information on TL and TL dynamics and to this end we ran a linear mixed-effects model with TL as dependent variable, age as covariate and bird ID as random effect. Gel ID was included as random effect to control for methodologically induced variance. With a log-likelihood ratio test, comparing models with and without bird ID, we tested for between-individual variation of TL. Further, including sex and its interaction with age in the model, we tested for differences in TL and its dynamics between females and males.

Second, we ran a series of univariate and bivariate ‘animal models’ (Wilson et al., 2010) using early-life TL data and pedigree information holding various degrees of kinship to partition the total phenotypic variance in early-life TL (*V*_*P*_) measured at the ages of 4 or 29 days into its additive genetic (*V*_*A*_), residual (*V*_*R*_) and varying other components of potential relevance: Genetic mother ID (V_gM_) estimates the variance in TL among nestlings from different mothers over and above the variance that is attributable to additive genetic effects. The TL in offspring from the same genetic mother may be more similar as compared to offspring from other mothers due to mother-specific effects, if e.g. egg content affects offspring TL (Haussmann, Longenecker, Marchetto, Juliano, & Bowden, 2012). Similarly, genetic father ID (*V*_*gF*_) and foster mother or father ID could affect TL of nestlings by parental care effects (e.g. Vedder, Verhulst, Zuidersma, & Bouwhuis, 2018). Birth year (*V*_*Y*_) was added to account for environmental variability between years that could affect nestling TL. Brood ID (*V*_*B*_) estimates the variance that can be attributed to the shared environment of brood mates. As the composition of the brood (number and identity of nestlings) changed after the experimental manipulation at the age of 4 days, individuals have a different brood ID pre- and post-manipulation (B1 and B2 respectively). To account for measurement differences between gels, we always included gel ID as random effect (*V*_*gel*_). As fixed effect we included father age (Bauch et al., 2019). Sex as fixed effect was non-significant and therefore not included in the final models presented. As we measured TL for the entire brood when the oldest nestling was 4 days old, and jackdaws hatch asynchronously, we tested whether the exact age of the nestling had an effect on TL. This was however not the case and therefore this predictor was also not included in the models presented. In the bivariate model with TL measured at both nestling ages we fitted brood ID until the age of 4 days (only genetic siblings and natural brood size, B1) for TL at age 4 days and brood ID of the following 25 days (genetic and non-genetic siblings, manipulated brood size, B2) for TL at age 29 days. The bivariate model was not improved if parent ID, birth year or gel ID were fitted separately for the two TL measurements. Statistical significance of random effects was calculated using likelihood ratio tests, comparing models with and without the specific random effect.

We calculated narrow sense heritability (*h*^*2*^) estimates for TL:

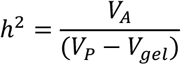

excluding variation due to gel effects (*V*_*gel*_). For a more comprehensive understanding of the *h*^2^- value, which can be affected by fixed effects, we re-ran the main model with random effects only (Wilson, 2008).

The genetic correlation *r*_G_ between TL at the ages 4 and 29 days was calculated as

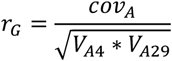

where cov_A_ is the additive genetic covariance between the repeated TL measurements and V_A_ is the additive genetic variance at age 4 or 29 days.

Accordingly, the phenotypic correlation *r*_P_ was calculated as

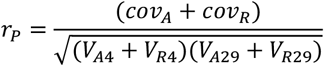

with cov_R_ as the residual covariance between the repeated TLs and V_R_ as the residual variance at age 4 or 29 days.

For TL at the two time points separately, the coefficient of biological variance of TL was calculated as

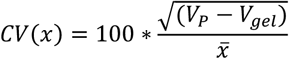

where x is TL and 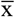 its mean, V_P_ is the total phenotypic variance from which *V*_gel_, the variance explained by gel differences, is subtracted (Houle, 1992).

The coefficient of additive genetic variance of TL at both time points was calculated as:

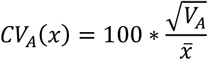

where x is TL and 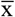 its mean and V_A_ is the additive genetic variance calculated in the animal model (Houle, 1992).

The relative evolvability I_A_ of TL at both time points was calculated as:

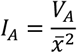

(Houle, 1992); multiplied by 100 to express it as a percentage.

Third, for comparison with the animal model results, we applied parent-offspring regressions using linear mixed-effects models of early-life TL (age 4 days) and the change in TL during the nestling period (between age 4 and 29 days) between parents and their offspring. We ran these analyses for mothers and fathers separately as data on both parents simultaneously were available in too few cases. When regressing offspring TL on father TL we ran an additional model that included father age as fixed effect, as TL of consecutive offspring declines as fathers age (Bauch et al., 2019). Additionally, we calculated a father-offspring regression with father TL measured in the year of offspring conception. As the dataset contains TL or TL change of offspring of the same parents in multiple years and TL was analysed on different gels, we added parent ID, brood ID, birth year and gel ID as random effects.

All statistics were run in R (R Development Core Team) using ASREML-R software (versions 3 and 4, VSN International Ltd.) and R-packages LME4 (Bates, Mächler, Bolker, & Walker, 2015) and LMERTEST (Kuznetsova, Brockhoff, & Christensen, 2017).

## Results

### Descriptive statistics

TL at the age of 4 days was on average 7039 bp (SD = 591 bp, n=715). Telomeres shortened within individuals over the measurement interval of 25 days by on average 260 bp (SD=171bp; t=-33.18, p<0.001, n=474). TL differed significantly between individuals (χ^2^=986.17; p<0.001), and was highly correlated within-individuals at the ages of 4 and 29 days (r=0.96, n=474, p<0.001, Fig. 1). The TL repeatability, corrected for age and gel effects, was 97% (F_474,2_=32.7, p=0.030; Lessells & Boag, 1987). The sexes did not differ in TL, neither at the age of 4 days (average TL±SD; females: 7036±603 [bp], n=357; males: 7031±581 [bp], n=339; F_1,659.84_=0.52, p=0.47), nor 25 days later (females: 6717±611 [bp], n=235; males: 6748±579 [bp], n=234; F_1,449.75_=1.35, p=0.25) and consequently also not in telomere loss (age*sex: t=0.59, p=0.55).

**Fig. 1.**
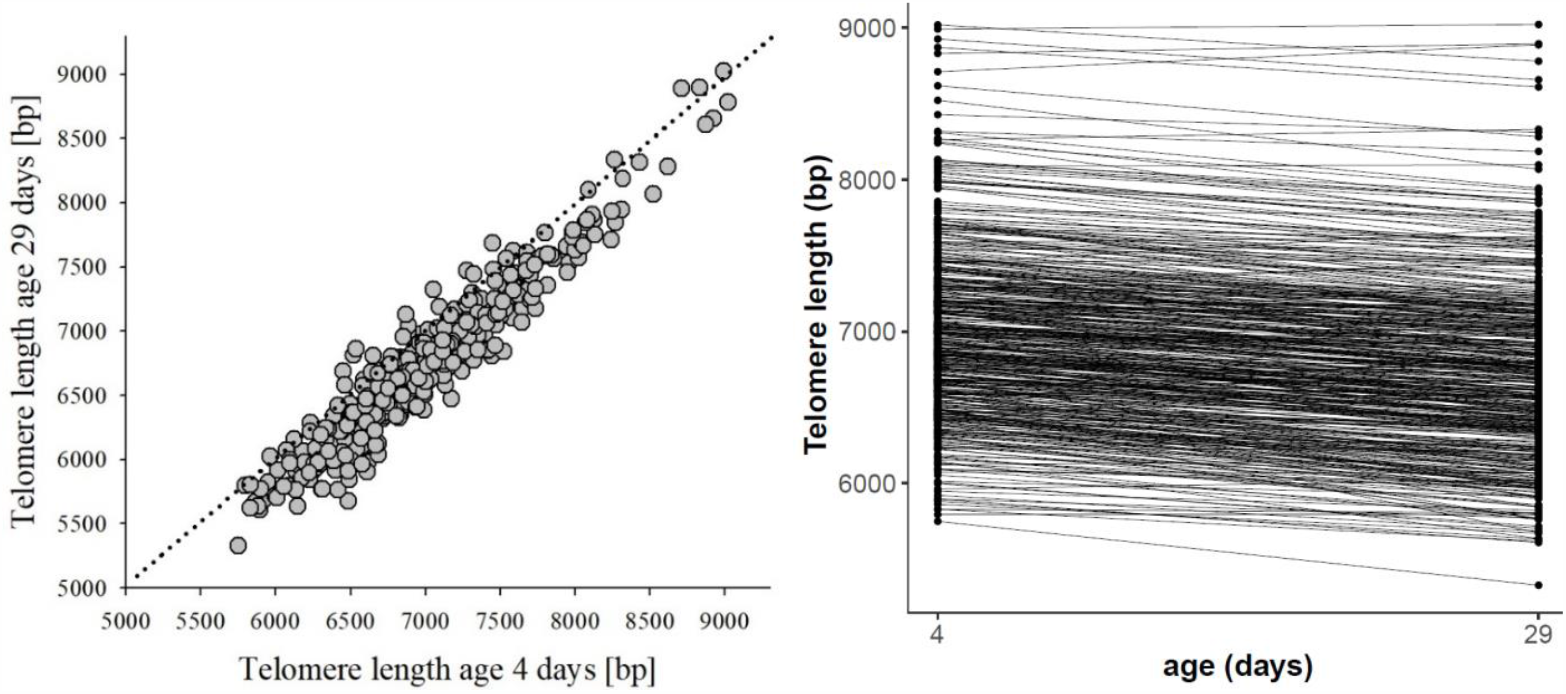
Repeated telomere length data within individuals (n=474). **(A)** Telomere length at 29 days of age plotted against telomere length at 4 days of age in jackdaw nestlings (r=0.96). The dashed line represents x=y and hence the perpendicular distance below this line reflects the telomere shortening. **(B)** Telomere length at the ages 4 and 29 days, where lines connect repeated data of the same individual. Average telomere loss over 25 days is 260 bp.

### Animal model analyses

The uni- and bivariate animal model analyses revealed that the major part of the variation in TL at the age of 4 and 29 days was explained by additive genetic effects (Tables 1, S2-S5). The narrow sense heritability estimates for TL measured at the ages of 4 and 29 days, derived from the bivariate model, were *h*^2^=0.71 and 0.77, respectively (Table 1, Fig. 2). Heritability estimates based on respective univariate models, thus not taking the relation of repeatedly measured TL within individuals into account, were *h*^2^=0.63 (lower) for TL measured at the age of 4 days (Table S2) and *h*^2^=0.88 (higher) for TL measured at the age of 29 days (Table S5). Including paternal age as a fixed effect in the model did not significantly lower the heritability estimate (*h*^2^=0.63 vs. 0.67, for TL measured at 4 days of age, Tables S2, S3).

**Table 1.**
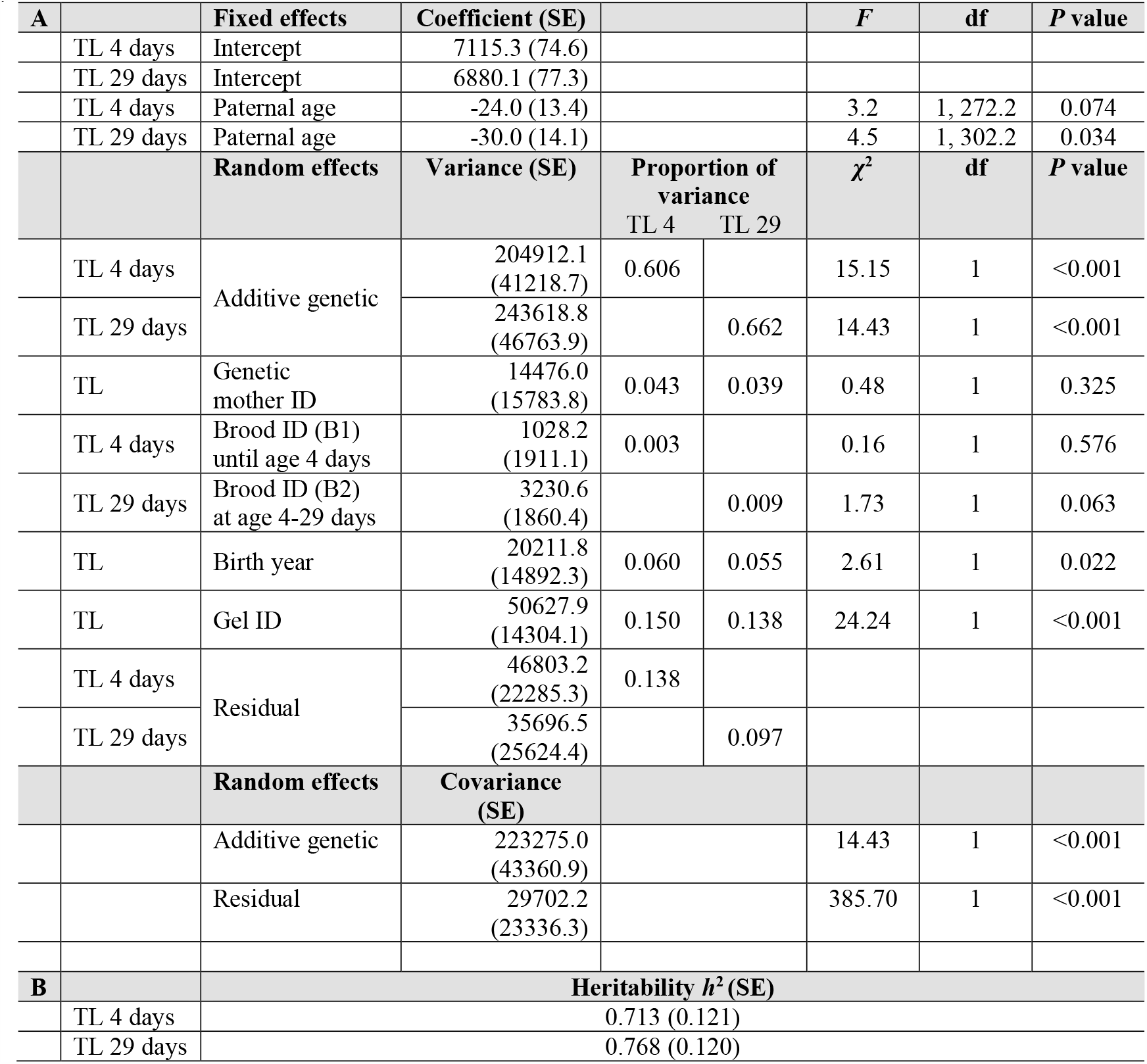
Results from a bivariate animal model analysis on telomere length measured at the ages 4 days in 715 individuals and again at 29 days in a subset of 474 individuals. **(A)** Fixed effects, variance components and covariances of telomere length. Note that brood ID prior to TL measurement at the ages 4 and 29 days differed due to experimental manipulation and accordingly the respective brood ID is fitted for the TL. Consequently, variance due to additive genetic effects and residual variance are also estimated separately, leading to different proportions of variance for TL at the age of 4 and 29 days. **(B)** Heritability estimates for telomere length.

**Fig. 2.**
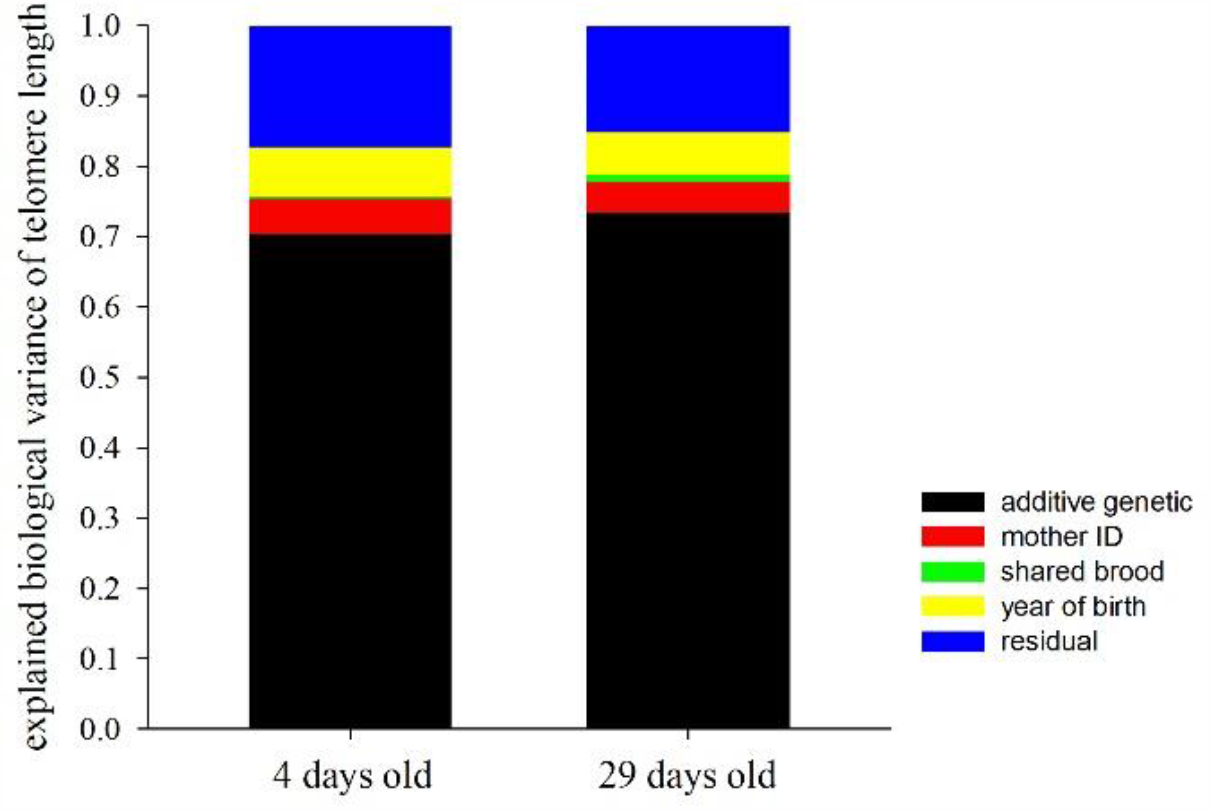
Components explaining biological variance of telomere length among individuals measured at the ages 4 days and 29 days derived from a bivariate animal model (Table 1). Additive genetic effects explained the majority of variance, while variance due to environmental effects was low and non-significant except for birth year. Components from bottom to top: additive genetic, mother ID, brood ID (note that it is hardly visible), birth year, residual.

Birth year was the only environmental factor that explained a significant part of the variance in early-life TL (Table 1), indicating resemblance of TL between chicks within cohorts. Genetic mother ID explained only very small, non-significant fractions of the variance in TL at the age of 4 and 29 days (Tables 1, S2, S3). Father ID in the model could not be estimated (Tables 1, S2-S5). Foster parent ID did not explain any significant TL variation. Brood ID explained little variation at both sampling ages (Tables 1, S2-S5).

TLs at the ages of 4 and 29 days were strongly genetically correlated (*r*_*G*_=0.999, SE: 0.006, p<0.001) and the additive genetic variances for the two TLs did not differ (p=0.61). Thus, there is no statistical support for a genotype by age interaction, and hence there is no evidence for additive genetic effects explaining between individual differences in TL change over this period. The coefficient of variation of phenotypic variance was 7.62% or 8.37% for TL at the ages of 4 days or 29 days, respectively. Accordingly, the coefficient of additive genetic variance was 6.43% or 7.34%. This resulted in estimates of the evolvability of TL of 0.41% and 0.54% for TL at the ages of 4 and 29 days.

### Parent-offspring regressions

TLs of mothers and their offspring, all measured at the age of 4 days, were significantly positively related (*β*=0.42±0.11), amounting to a to TL heritability estimate of *h*^*2*^=0.84 (Table 2A, Fig 3A). The same relationship for TLs of fathers and their offspring was non-significant, whether paternal age was included in the model (*β*=0.10±0.11, Table 2B) or not (*β*=0.18±0.10, Table S6; Fig. 3B). Older fathers produced chicks with shorter TL as reflected in a significant negative paternal age effect. The slope of the mother-offspring regression was more than twice as steep as the slope derived from the father-offspring regression (Table 2). In contrast, paternal TL measured at the age of offspring conception was significantly positively related to offspring early-life TL (*β*=0.31±0.07; Table S7), amounting to a TL heritability estimate of *h*^*2*^=0.61.

**Table 2.** Parent-offspring regressions. Comparing telomere length measured at the age of 4 days (in bp) of **(A)** mothers and their offspring (n = 113 individual offspring of 31 mothers) and **(B)** fathers and their offspring (n = 111 individual offspring of 28 fathers).

| **A** |  |  |  |  |  |
| --- | --- | --- | --- | --- | --- |
| **Fixed effects** | **Coefficient (SE)** |  | **F** | **df** | **P value** |
| Intercept | 4283.6 (781.6) |  |  |  |  |
| Mother's TL<br>(age 4 days) | 0.418 (0.106) |  | 15.44 | 1, 16.4 | 0.001 |
| **Random effects** | **Variance (SE)** | **Proportion of variance (SE)** | **$\chi^2$** | **df** | **P value** |
| Mother ID | 71974.6 (41762.7) | 0.205 (0.110) | 3.940 | 1 | 0.047 |
| Brood ID (B1)<sup>a</sup> | - | - | - | - | - |
| Birth year | 92126.7 (66701.9) | 0.263 (0.148) | 4.763 | 1 | 0.029 |
| Gel ID | 14888.4 (23115.2) | 0.042 (0.067) | 0.521 | 1 | 0.471 |
| Residual | 171639.6 (31312.4) | 0.490 (0.125) |  |  |  |
| **B** |  |  |  |  |  |
| **Fixed effects** | **Coefficient (SE)** |  | **F** | **df** | **P value** |
| Intercept | 6865.4 (830.0) |  |  |  |  |
| Father's TL<br>(age 4 days) | 0.102 (0.110) |  | 0.861 | 1, 25.6 | 0.362 |
| Father's age at<br>conception | -88.6 (38.2) |  | 5.386 | 1, 20.1 | 0.031 |
| **Random effects** | **Variance (SE)** | **Proportion of variance (SE)** | **$\chi^2$** | **df** | **P value** |
| Father ID | 62097.5 (46496.8) | 0.200 (0.134) | 1.948 | 1 | 0.163 |
| Brood ID (B1) | 27415.8 (45317.5) | 0.088 (0.147) | 0.297 | 1 | 0.586 |
| Birth year | 11885.5 (22898.4) | 0.038 (0.072) | 0.446 | 1 | 0.504 |
| Gel ID | 7893.1 (23205.1) | 0.025 (0.075) | 0.134 | 1 | 0.715 |
| Residual | 200755.8 (40829.6) | 0.647 (0.137) |  |  |  |

<sup>a</sup> Brood ID was not included in the final model as it could not be estimated.

**Fig. 3.**
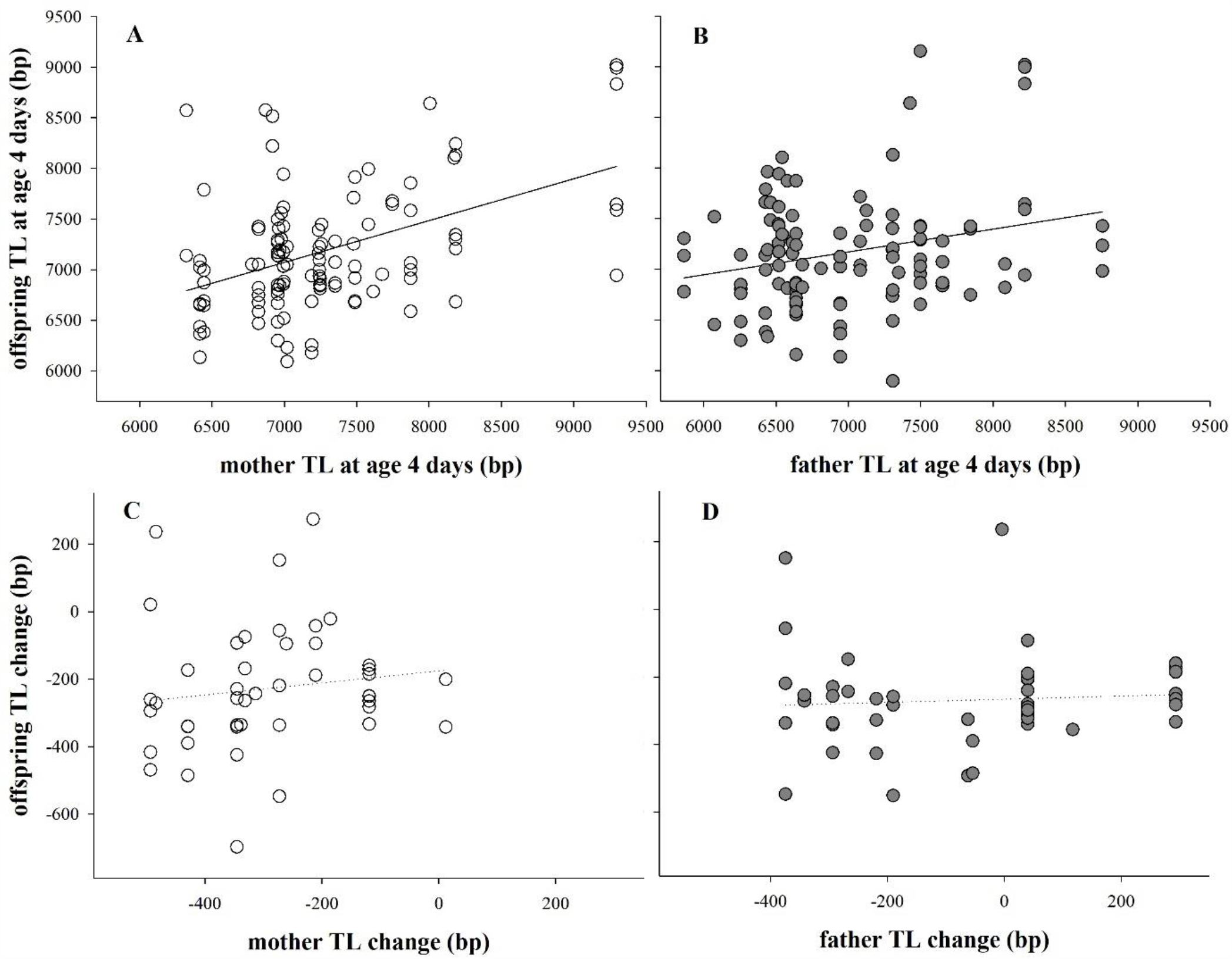
Parent-offspring regressions on telomere length and its change. Telomere length measured at the age of 4 days in **(A)** mothers (n=31) and offspring (n=113) and **(B)** fathers (n=28) and offspring (n=111). For statistics see table 2. Change in telomere length between the ages of 4 and 29 days in **(C)** mothers (n=14) and offspring (n=45) and **(D)** fathers (n=12) and offspring (n=45). For statistics see table S8.

TL change from age 4 to 29 days was neither significantly correlated between mother and offspring (*β*=0.18±0.24), nor between father and offspring (*β*=0.04±0.17; Table S8, Fig. 3C-D). Thus, parent-offspring regressions confirm the lack of statistical support for heritable variation in telomere shortening.

## Discussion

We found that TL was highly heritable. Our estimate of TL heritability in jackdaws is similar to estimates reported by other studies on free-living bird species such as terns and swallows (range *h*^2^=0.63-0.81; Table 3), but substantially higher than in studies on penguins, dippers, warblers and flycatchers (range *h*^2^=0.0-0.48; Table 3). One striking difference between those studies within the same taxonomic group (Aves) is, that they used different telomere measurement techniques, TRF and qPCR, respectively. Interestingly, TRF- and qPCR-based studies also report differences in TL repeatability, being high and low, respectively (e.g. this study; Benetos et al., 2019; Bichet et al., 2020 vs. Foley et al. 2020; Sparks et al, 2020; van Lieshout et al., 2020; meta-analysis: Kärkkäinen et al. in prep). This is of relevance as TL repeatability sets, with exceptions, the upper limit to heritability (Dohm, 2002). The difference in heritability and repeatability estimates between TRF- and qPCR-based studies may be due to chance, given that the number of studies is still small, but may also be caused by real differences between the methods. With respect to the latter possibility, two main differences between the method of TRF and qPCR that may contribute to variation in repeatability and heritability estimates are that (i) qPCR-based measurements include all telomeric sequences within the genome, thus terminally located and ITS, which are excluded when applying TRF to non-denatured DNA, and (ii) measurement reliability tends to be lower when using qPCR (Horn, Robertson, & Gemmell, 2010; Morinha, Magalhães, & Blanco, 2020; Nussey et al. 2014). As TRF requires high DNA integrity to work, samples for those studies are usually taken and stored explicitly for TL measurement, while various storage methods have been reported for samples processed with qPCR (Eastwood, Mulder, Verhulst, & Peters, 2017). Thus, sample quality may be less variable in TRF-based studies. As for the potential influence of ITS on TL, their abundance varies between and within species (Delany, Krupkin, & Miller, 2000; Foote et al. 2013; Meyne et al. 1990; Olsson et al. 2011). However, for the inclusion of ITS to explain the lower repeatability and heritability estimates when using qPCR, ITS would have to be more susceptible to environmental effects than terminal telomeres, resulting in a higher absolute change in the total number of telomere sequence repeats in the genome. There is at present little indication that this is a likely scenario. We therefore consider measurement error, either caused by sample quality or the measurement process, the most parsimonious explanation for the difference in repeatability and heritability estimates between qPCR- and TRF-based studies.

**Table 3.** Overview of the studies estimating heritability (*h*^*2*^) of telomere length (TL) in non-human vertebrates.

| | Species | age TL measured | cross-foster | controlled environment | father age effect | statistical analysis | $h^2$ (95% CI) | *n* | cell type | lab method | reference |
| --- | --- | --- | --- | --- | --- | --- | --- | --- | --- | --- | --- |
| birds | king penguin<br>*Aptenodytes patagonicus* | offspring 10 days, parents at conception | no | no | not reported | mid-parent-offspring<br>mother-offspring | **0.2 (-0.02-0.42)**<br>**0.2 (0.01-0.39)** | 53<br>53 | erythrocytes | qPCR | Reichert et al. 2015 |
|  | white-throated dipper<br>*Cinclus cinclus* | 7-17 days | no | no<br>brood ID cohort | not reported | mother-mid-offspring<br>father-mid-offspring<br>animal model | 0.44 (0.048-0.83)<br>0.08 (-0.35-0.51)<br>**0.038 (-0.10-0.17)** | 59<br>59<br>177 | erythrocytes | qPCR | Becker et al. 2015 |
|  | great reed warbler<br>*Acrocephalus arundinaceus* | 8-10 days | no | no<br>brood ID<br>mother ID | no, but maternal age effect | mother-mid-offspr.<br>father-mid-offspring<br>animal model | 1.08<br>n.s.<br>**0.48 (0.24-0.72)** | 17<br>19<br>193 | erythrocytes | qPCR | Asghar et al. 2015 |
|  | collared flycatcher<br>*Ficedula albicollis* | 12 days | yes | brood triplets | not reported | animal model (full siblings) | **0.09 (-0.04-0.15)** | 359 | erythrocytes | qPCR | Voillemot et al. 2012 |
|  | Seychelles warbler<br>*Acrocephalus sechellensis* | all ages | no | no<br>mother & father ID, sample & birth year, territory | yes | mother-offspring<br>father-offspring<br>animal model | n.s.<br>n.s.<br>**0.031 (<0.001-0.067)** | 303<br>284<br>1318 (2664 samples) | erythrocytes | qPCR | Sparks et al. 2020 |
|  | kakapo<br>*Strigops habroptilus* | 1-35 years | no | no | not reported | mother-offspring<br>father-offspring | **0.84**<br>n.s. | 29<br>26 | erythrocytes | TRF<sup>a</sup> | Horn et al. 2011 |
|  | tree swallow<br>*Tachycineta bicolor* | offspr. 12 days, parents at conception | yes | brood size | no | mid-parent-offspring | **0.81** | 122 | erythrocytes | TRF | Belmaker et al. 2019 |
|  | zebra finch<br>*Taeniopygia guttata* | 9-636 days | yes | birth nest<br>rear nest<br>mother ID<br>father ID | not reported | animal model | **0.999 (0.87-1)** | 125 | erythrocytes | TRF | Atema et al. 2015 |
|  | common tern<br>*Sterna hirundo* | 2-24 years | no | permanent environment | yes | animal model | **0.63 (0.50-0.73)** | 387 (619 samples) | erythrocytes | TRF | Vedder et al. under review |
|  | jackdaw<br>*Corvus monedula* | 4 days + 29 days<br>fathers at conception (ac) | yes | brood ID<br>mother ID<br>father ID<br>birth year | yes | mother - offspring<br>father - offspring<br>father ac - offspring<br>biv. animal model | 0.84 (0.42-1)<br>0.20 (-0.23-0.63)<br>0.61 (0.32-0.90)<br>**0.71 (0.48-0.95)**<br>**0.77 (0.53-1.0)** | 113<br>111<br>352<br>715 (1189 samples) | erythrocytes | TRF | this study |

| table 3 continued |  |  |  |  |  |  |  |  |  |  |  |
| --- | --- | --- | --- | --- | --- | --- | --- | --- | --- | --- | --- |
| | Species | age TL measured | cross-foster | controlled environment | father age effect | statistical analysis | $h^2$ (95% CI) | $n$ | cell type | lab method | reference |
| reptiles | sand lizard<br>*Lacerta agilis* | hatchlings, known-age adults | no | no | yes | mother-daughter, father-son | **0.52 (0.09-0.95)**<br>**1.23 (0.80-1.66)** | 55<br>40 | erythrocytes | TRF<sup>a</sup> | Olsson et al. 2011 |
| fish | nine-spined sticklebacks<br>*Pungitius pungitius* | 122 days | yes<sup>b</sup> | temperature family ID | not tested | animal models (full siblings of 9 families) | **0.31 (0.05–0.97)**<br>**to 0.47(0.17–0.91)** | 83; 213 | brain tissue | qPCR<sup>c</sup> | Noreikiene et al. 2017 |
| mammals | greater mouse-eared bat<br>*Myotis myotis* | 0-7+ years | no | birth year<br>sample year<br>roost<br>temperature<br>rainfall | not tested | animal model | **0.01 (0.00-0.04)**<br>**to 0.06 (0.02-0.11)** | 174 (504 samples) | wing biopsy | qPCR | Foley et al. 2020 |
|  | Holstein Friesian dairy cattle | all at birth + later in life | no | birth year<br>genetic group | not tested | random regression model incl. pedigree information | **0.36 to 0.47 (SE 0.05-0.08)** | 308 females (1328 samples) | leucocytes | qPCR | Seeker et al. 2018 |
|  | badger *Meles meles* | ≤29 months<br>all ages | no | mother ID, father ID, birth year, sample year, social group | no | animal model | **<0.001 (<0.001-0.026)**<br>**<0.001 (<0.001-0.043)** | 556 juveniles (837 samples)<br>612 (1248 samples) | leucocytes | qPCR | van Lieshout et al. 2020 |

<sup>a</sup>TRF including interstitial telomeric repeats (Southern Blot); <sup>b</sup> reared individually after hatching; <sup>c</sup> no ITS in the genome

Insufficient statistical control for shared environment effects has been suggested as another source for variation in TL heritability estimates between studies (Dugdale & Richardson, 2018). However, TL heritability estimates in our study were high, independent of statistical approach, due to small early-life parental and other environmental effects (Tables 1a, S1, and 2). Among the tested environmental components, birth year effects explained a significant but small part of the variation in TL in our study population (6 and 5.5% at ages 4 and 29 days respectively, Table 2). Birth cohort effects on TL have also been identified in other free-living vertebrates such as white-throated dippers *Cinclus cinclus* (Becker et al., 2015) and Soay sheep *Ovis aries* (Fairlie et al., 2016). However, the proportion of TL variance explained by birth year differed strongly between studies (6% in our study vs. 46% in Becker et al., 2015). The cause of variation in TL between birth cohorts remains to be identified.

The correlations of early-life TL of mother and offspring were stronger than between father and offspring and hence suggest TL to be mainly maternally inherited. As previously shown, older fathers produced offspring with shorter TL (paternal age effect) in this (Bauch et al. 2019, Table 2b), suggesting an epigenetic effect via declining TL in ageing fathers that is transferred to offspring via sperm. Correspondingly, father TL at the age of conception and offspring TL at the age of 4 days were significantly positively related in our study. Thus, taking into account paternal epigenetic inheritance, our results support a similar quantitative contribution of maternal and paternal TL to TL inheritance.

We did not find any statistical support for a significant heritable component in the rate of TL shortening. TL at the two ages was strongly genetically correlated, thus not supporting a genetic variation of TL among genotypes with age. Furthermore, parent-offspring regressions revealed no significant correlations in TL shortening, although the non-significant heritability estimates (mother-offspring *h*^*2*^=0.36±0.47, father-offspring *h*^*2*^=0.08±0.35) were on average (0.22) close to the only one other study of which we are aware that investigated heritability of TL dynamics, which reported *h*^*2*^=0.28 based on adult human twins (Hjelmborg et al., 2015). Whether genotype-dependent effects on TL variation change with advancing age, as (weakly) suggested in cattle (Seeker et al., 2018), remains to be tested. However, to detect such a (statistically significant) effect in adulthood, when telomere shortening is generally lower than early in life (Salomons et al., 2009), will require a much larger sample size and/or very long-term investigations.

It is tempting to suppose that a high heritability of TL indicates that TL would change fast in response to directional selection. However, we found the coefficient of additive genetic variance of TL to be low (6.43% or 7.34%) and consequently its potential to evolve is low, too. TL evolvability was also low in field crickets (Boonekamp et al., 2020a), which fits the expectations for a trait related to fitness (Houle et al. 1992), where high selection pressures and trait canalisation are expected (Stearns, Kaiser, & Kawecki, 1995; Vedder, Verhulst, Bauch, & Bouwhuis, 2018). But more studies that provide coefficients of additive genetic variation for comparison among populations and taxa are required to gain a more comprehensive picture.

### Ethics

Data were collected under license of the animal experimentation committee of the University of Groningen (license numbers 4071, 5871, 6832A). License was awarded in accordance with the Dutch national law on animal experimentation (“Wet op de dierproeven”).

## Acknowledgements

We thank Martijn Salomons, the late Cor Dijkstra and the many helpers involved in data and blood sample collection in this long-term project, as well as Erica Zuidersma, who contributed to the analyses of TLs. CB was supported by a DFG research fellowship (BA 5422/1-1) and JJB by an NWO grant (823.01.009) awarded to SV.

## Data accessibility

The data that support the findings of this study are available from the Dryad Digital Repository: http://dx.doi.org/…*to be completed upon acceptance for publication*

## Author contributions

Project design and organisation by SV; fieldwork by JJB, SV, CB; labwork by EM, CB; data analysis by CB, PK, JJB; manuscript first version by CB; all authors contributed to the final version.

## References

Angelier, F., Costantini, D., Blévin, P., & Chastel, O. (2018) Do glucocorticoids mediate the link between environmental conditions and telomere dynamics in wild vertebrates? A review. General and Comparative Endocrinology, 256, 99–111. doi:10.1016/j.ygcen.2017.07.007

Asghar, M., Bensch, S., Tarka, M., Hansson, B., & Hasselquist, D. (2015) Maternal and genetic factors determine early life telomere length. Proceedings of the Royal Society B: Biological Sciences, 282, 20142263. doi:10.1098/rspb.2014.2263

Atema, E., Mulder, E., Dugdale, H. L., Briga, M., van Noordwijk, A. J., & Verhulst S. (2015) Heritability of telomere length in the Zebra Finch. Journal of Ornithology, 156, 1113–1123. doi:10.1007/s10336-015-1212-7

Bates, D., Mächler, M., Bolker, B., & Walker, S. (2015) Fitting Linear Mixed-Effects Models Using lme4. Journal of Statistical Software, 67, 1–48. doi:10.18637/jss.v067.i01

Bauch, C., Becker, P. H., & Verhulst, S. (2013) Telomere length reflects phenotypic quality and costs of reproduction in a long-lived seabird. Proceedings of the Royal Society of London B: Biological Sciences, 280, 20122540. doi:10.1098/rspb.2012.2540

Bauch, C., Boonekamp, J. J., Korsten, P., Mulder, E., & Verhulst, S. (2019) Epigenetic inheritance of telomere length in wild birds. PLoS Genetics, 15, e1007827. doi:10.1371/journal.pgen.1007827

[dataset] Bauch, C., Korsten, P., Boonekamp, J. J., Mulder, E., & Verhulst S. (2020) Data from: High heritability of telomere length, but low evolvability, and no significant heritability of telomere shortening. Dryad Digital Repository, doi … filled upon acceptance

Becker, P. J. J., Reichert, S., Zahn, S., Hegelbach, J., Massemin, S., Keller, L. F.,… Criscuolo, F. (2015) Mother-offspring and nest-mate resemblance but no heritability in early-life telomere length in white-throated dippers. Proceedings of the Royal Society B: Biological Sciences, 282, 20142924. doi:10.1098/rspb.2014.2924

Belmaker, A., Hallinger, K. K., Glynn, R. A., Winkler, D. W., & Haussmann, M. F. (2019) The environmental and genetic determinants of chick telomere length in Tree Swallows (Tachycineta bicolor). Ecology and Evolution, 9, 8175–8186. doi:10.1002/ece3.5386

Benetos, A., Verhulst, S., Labat, C., Lai, T.-P., Girerd, N., Toupance, S.,… Aviv, A. (2019) Telomere length tracking in children and their parents: implications for adult onset diseases. The FASEB Journal, 33, 14248–14253. doi:10.1096/fj.201901275R

Bichet, C., Bouwhuis, S., Bauch, C., Verhulst, S., Becker, P. H., & Vedder, O. (2020) Telomere length is repeatable, shortens with age and reproductive success, and predicts remaining lifespan in a long-lived seabird. Molecular Ecology, 29, 429–441. doi:10.1111/mec.15331

Blackburn, E. H. (1991) Structure and function of telomeres. Nature, 350, 569–573. doi:10.1146/annurev.genet.23.1.579

Boonekamp, J. J., Bauch, C., & Verhulst, S. (2020b) Experimentally increased brood size accelerates actuarial senescence and increases subsequent reproductive effort in a wild bird population. Journal of Animal Ecology, 89, 1395–1407. doi:10.1111/1365-2656.13186

Boonekamp, J. J., Mulder, G. A., Salomons, H. M., Dijkstra, C., & Verhulst, S. (2014) Nestling telomere shortening, but not telomere length, reflects developmental stress and predicts survival in wild birds. Proceedings of the Royal Society B: Biological Sciences, 281, 20133287. doi:10.1098/rspb.2013.3287

Boonekamp, J. J., Rodriguez-Munoz, R., Hopwood, P., Zuidersma, E., Mulder, E., Wilson, A., Tregenza T. (2020a) Telomere length is highly heritable and independent of growth rate manipulated by temperature in field crickets. bioRxiv. doi:10.1101/2020.05.29.123216

Boonekamp, J. J., Simons, M. J. P., Hemerik, L., & Verhulst, S. (2013) Telomere length behaves as biomarker of somatic redundancy rather than biological age. Aging Cell, 12, 330–332. doi:10.1111/acel.12050

Broer, L., Codd, V., Nyholt, D. R., Deelen, J., Mangino, M., Willemsen, G,… Boomsma D. I. (2013) Meta-analysis of telomere length in 19 713 subjects reveals high heritability, stronger maternal inheritance and a paternal age effect. European Journal of Human Genetics, 21, 1163–1168. doi:10.1038/ejhg.2012.303

Chan, S. R. W. L., & Blackburn, E. H. (2004) Telomeres and Telomerase. Philosophical Transactions of the Royal Society London B, 359, 109–121. doi:10.1098/rstb.2003.1370

de Lange, T. (2005) Shelterin: the protein complex that shapes and safeguards human telomeres. Genes & Development, 19, 2100–2110. doi:10.1101/gad.1346005

Delany, M. E., Krupkin, A. B., & Miller, M. M. (2000) Organization of telomere sequences in birds: evidence for arrays of extreme length and for in vivo shortening. Cytogenetics and Cell Genetics, 90, 139–145.

Dohm, M. R. (2002) Repeatability Estimates Do Not Always Set an Upper Limit to Heritability. Functional Ecology, 16, 273–280. https://www.jstor.org/stable/826658

Dugdale, H. L., & Richardson, D. S. (2018) Heritability of telomere variation: it is all about the environment. Philosophical Transactions of the Royal Society B, 373, 20160450. doi:10.1098/rstb.2016.0450

Eastwood, J. R., Mulder, E., Verhulst, S., & Peters, A. (2017) Increasing the accuracy and precision of relative telomere length estimates by RT qPCR. Molecular Ecology Resources, 18, 68–78. doi:10.1111/1755-0998.12711

Eisenberg, D. T. A. (2019) Paternal age at conception effects on offspring telomere length across species – What explains the variability? PLoS Genetics, 15, e1007946. doi:10.1371/journal.pgen.1007946

Fairlie, J., Holland, R., Pilkington, J. G., Pemberton, J. M., Harrington, L., & Nussey, D. H. (2016) Lifelong leukocyte telomere dynamics and survival in a free-living mammal. Aging Cell, 15, 140–148. doi:10.1111/acel.12417

Falconer, D. S., & Mackay, T. F. C. (1996) Introduction to quantitative genetics (4th ed). Harlow, Essex.

Foley, N., Petit, E. J., Brazier, T., Finarelli, J. A., Hughes, G. M., Touzalin, F.,… Teeling, E. C. (2020) Drivers of longitudinal telomere dynamics in a long-lived bat species, Myotis myotis. Molecular Ecology, 29, 2963–2977. doi:10.1111/mec.15395

Foote, C. G., Vleck, D., & Vleck, C. M. (2013) Extent and variability of interstitial telomeric sequences and their effects on estimates of telomere length. Molecular Ecology Resources, 13, 417–428. doi:10.1111/1755-0998.12079

Griffiths, R., Double, M. C., Orr, K., & Dawson, R. J. G. (1998) A DNA test to sex most birds. Molecular Ecology, 7, 1071–1075.

Hall, M. E., Nasir, L., Daunt, F., Gault, E. A., Croxall, J. P., Wanless, S., & Monaghan, P. (2004) Telomere loss in relation to age and early environment in long-lived birds. Proceedings of the Royal Society of London B: Biological Sciences, 271, 1571–1576. doi:10.1098/rspb.2004.2768

Haussmann, M. F., Longenecker, A. S., Marchetto, N. M., Juliano, S. A., & Bowden, R. M. (2012) Embryonic exposure to corticosterone modifies the juvenile stress response, oxidative stress and telomere length. Proceedings of the Royal Society of London B: Biological Sciences, 279, 1447–1456. doi:10.1098/rspb.2011.1913

Henderson, I. G., Hart, P. J. B., & Burke, T. (2000) Strict monogamy in a semi-colonial passerine: the Jackdaw Corvus monedula. Journal of Avian Biology, 31, 177–182. doi:10.1034/j.1600-048X.2000.310209.x

Hjelmborg, J. B., Dalgård, C., Möller, S., Steenstrup, T., Kimura, M., Christensen, K.,… Aviv, A. (2015) The heritability of leucocyte telomere length dynamics. Journal of Medical Genetics, 52, 297–302. doi:10.1136/jmedgenet-2014-102736

Horn, T., Robertson, B. C., & Gemmell, N. J. (2010) The use of telomere length in ecology and evolutionary biology. Heredity, 105, 497–506. doi:10.1038/hdy.2010.113

Horn, T., Robertson, B. C., Will, M., Eason, D. K., Elliott, G. P., & Gemmell, N. J. (2011) Inheritance of telomere length in a bird. PLoS ONE, 6, e17199. doi:10.1371/journal.pone.0017199

Houle, D. (1992) Comparing evolvability and variability of quantitative traits. Genetics, 130, 195–204.

Kotrschal, A., Ilmonen, P., & Penn, D. J. (2007) Stress impacts telomere dynamics. Biology Letters, 3, 128–130. doi:10.1098/rsbl.2006.0594

Kruuk, L. E. B., & Hadfield, J. D. (2007) How to separate genetic and environmental causes of similarity between relatives. Journal of Evolutionary Biology, 20, 1890–1903. doi:10.1111/j.1420-9101.2007.01377.x

Kuznetsova, A., Brockhoff, P. B., & Christensen, R. H. B. (2017) lmerTest Package: Tests in Linear Mixed Effects Models. Journal of Statistical Software, 82, 1–26. Version 3.1-0. doi:10.18637/jss.v082.i13

Lessells, C. M., & Boag, P. T. (1987) Unrepeatable repeatabilities: a common mistake. The Auk, 104, 116–121. doi:10.2307/4087240

Liebers, D., & Peter, H.-U. (1998) Intraspecific interactions in jackdaws Corvus monedula: A field study combined with parentage analysis. Ardea, 86, 221–235.

McLennan, D., Amstrong, J. D., Stewart, D. C., McKelvey, S., Boner, W., Monaghan, P., & Metcalfe, N. M. (2016) Interactions between parental traits, environmental harshness and growth rate in determining telomere length in wild juvenile salmon. Molecular Ecology, 25, 5425–5438. doi:10.1111/mec.13857

Meyne, J., Baker, R. J., Hobart, H. H., Hsu, T. C., Ryder, O. A., Ward, O. G.,… Moyzis, R. K. (1990) Distribution of non-telomeric sites of the (TTAGGG)n telomeric sequence in vertebrate chromosomes. Chromosoma, 99, 3–10.

Monaghan, P., Eisenberg, D. T. A., Harrington, L., & Nussey, D. H. (Eds.). (2018) Understanding diversity in telomere dynamics [Special issue]. Philosophical Transactions of the Royal Society B Biological Sciences, 373, 1741.

Morinha, F., Magalhães, P., & Blanco, G. (2020) Standard guidelines for the publication of telomere qPCR results in evolutionary ecology. Molecular Ecology Resources, 20, 635–531 648. doi:10.1111/1755-0998.13152

Noguera, J. C., Metcalfe, N. B., & Monaghan, P. (2018) Experimental demonstration that offspring fathered by old males have shorter telomeres and reduced lifespans. Proceedings of the Royal Society B: Biological Sciences, 285, 20180268. doi:10.1098/rspb.2018.0268

Noreikiene, K., Kuparinen, A., & Merilä, J. (2017) Age at maturation has sex- and temperature-specific effects on telomere length in a fish. Oecologia, 184, 767–777. doi:10.1007/s00442-017-3913-5

Nussey, D. H., Baird, D., Barrett, E., Boner, W., Fairlie, J., Gemmell, N.,… Monaghan P (2014) Measuring telomere length and telomere dynamics in evolutionary biology and ecology. Methods in Ecology and Evolution, 5, 299–310. doi:10.1111/2041-210X.12161

Olsson, M., Pauliny, A., Wapstra, E., Uller, T., Schwartz, T., & Blomqvist, D. (2011) Sex differences in sand lizard telomere inheritance: paternal epigenetic effects increases telomere heritability and offspring survival. PLoS ONE, 6, e17473. doi:10.1371/journal.pone.0017473

O’Sullivan, R. J., & Karlseder, J. (2010) Telomeres: protecting chromosomes against genome instability. Nature Reviews: Molecular Cell Biology, 11, 171–181. doi:10.1038/nrm2848

R Core Team (2017) R: A language and environment for statistical computing. R Foundation for Statistical Computing, Vienna, Austria.

Reichert, S., Rojas, E. R., Zahn, S., Robin, J. P., Criscuolo, F., & Massemin, S. (2015) Maternal telomere length inheritance in the king penguin. Heredity, 114, 10–16. doi:10.1038/hdy.2014.60

Röell, A. (1978) Social behaviour of the jackdaw, Corvus monedula, in relation to its niche. Behaviour, 64, 1–124.

Salomons, H. M., Mulder, G. A., van de Zande, L., Haussmann, M. F., Linskens, M. H. K., & Verhulst, S. (2009) Telomere shortening and survival in free-living corvids. Proceedings of the Royal Society of London B: Biological Sciences, 276, 3157–3165. doi:10.1098/rspb.2009.0517

Seeker, L. A., Ilska, J. J., Psifidi, A., Wilbourn, R. V., Underwood, S. L., Fairlie, J.,… Banos, G. 2018 Longitudinal changes in telomere length and associated genetic parameters in dairy cattle analysed using random regression models. PLoS ONE, 13, e0192864. doi:10.1371/journal.pone.0192864

Sparks, A. M., Spurgin, L. G., van der Velde, M., Fairfield, E. A., Komdeur, J., Burke, T.,… Dugdale, H. L. (2020) Telomere heritability and parental age at conception effects in a wild avian population. EcoEvoRxiv. doi:10.32942/osf.io/eq2af

Spurgin, L. G., Bebbington, K., Fairfield, E. A., Hammers, M., Komdeur, J., Burke, T.,… Richardson, D. S. (2018) Spatio-temporal variation in lifelong telomere dynamics in a long-term ecological study. Journal of Animal Ecology, 87, 187–198. doi:10.1111/1365-2656.12741

Stearns, S. C., Kaiser, M., & Kawecki, T. J. (1995) The differential genetic and environmental canalization of fitness components in Drosophila melanogaster. Journal of Evolutionary Biology, 8, 539–557.

van Lieshout, S. H. J., Sparks, A. M., Bretman, A., Newman, C., Buesching, C. D., Burke, T., Dugdale, H. L. (2020) Estimation of environmental, genetic and parental age at conception effects on telomere length in a wild mammal. Journal of Evolutionary Biology. doi:10.1111/JEB.13728

Vedder, O., Verhulst, S., Bauch, C., & Bouwhuis, S. (2017) Telomere attrition and growth: a life-history framework and case study in common terns. Journal of Evolutionary Biology, 30, 1409–1419. doi:10.1111/jeb.13119

Vedder, O., Verhulst, S., Zuidersma, E., & Bouwhuis, S. (2018) Embryonic growth rate affects telomere attrition: an experiment in a wild bird. Journal of Experimental Biology, 221, jeb181586. doi:10.1242/jeb.181586

Voillemot, M., Hine, K., Zahn, S., Criscuolo, F., Gustafsson, L., Doligez, B., & Bize, P. (2012) Effects of brood size manipulation and common origin on phenotype and telomere length in nestling collared flycatchers. BMC Ecology, 12, 17.

Wilbourn, R. V., Moatt, J. P., Froy, H., Walling, C. A., Nussey, D. H., & Boonekamp, J. J. (2018) The relationship between telomere length and mortality risk in non-model vertebrate systems: a meta-analysis. Philosophical Transactions of the Royal Society B, 373, 20160447. doi:10.1098/rstb.2016.0447

Wilson, A. J. (2008) Why h2does not always equal VA/VP? Journal of Evolutionary Biology, 21, 647–650. doi:10.1111/j.1420-9101.2008.01500.x

Wilson, A. J., Réale, D., Clements, M. N., Morrissey, M. M., Postma, E., Walling, C. A.,… Nussey DH. (2010) An ecologist’s guide to the animal model. Journal of Animal Ecology, 79, 13–26. doi:10.1111/j.1365-2656.2009.01639.x

Young, R. C., Barger, C. P., Dorresteijn, I., Haussmann, M. F., & Kitaysky, A. S. (2013) Telomere length and environmental conditions predict stress levels but not parental investment in a long-lived seabird. Marine Ecology Progress Series, 556, 251–259. doi:10.3354/meps11864

