## Supplementary Material for "High heritability of telomere length, but low evolvability, and no significant heritability of telomere shortening in wild jackdaws"

for

Content:

- Telomere restriction fragment analysis method page 2
- Pedigree information (Table S1, Fig. S1) pages 3-4
- Animal models (Tables S2-S5) pages 5-6
- Parent-offspring regressions (Tables S6-S8) pages 7-8
- Reference page 8

#### *Telomere restriction fragment analysis method*

For the telomere restriction fragment analysis we removed the glycerol buffer, washed the cells and isolated DNA from 5 µl of erythrocytes using CHEF Genomic DNA Plug kit (Bio-Rad, Hercules, CA, USA). Cells in the agarose plugs were digested overnight with *Proteinase K* at 50°C. DNA in half of a plug per sample was restricted simultaneously with *Hind*III (60 U), *Hinf*I (30 U) and *Msp*I (60 U) for ~18 h in NEB2 buffer (New England Biolabs Inc., Beverly, MA, USA). The restricted DNA was then separated by pulsed-field gel electrophoresis in a 0.8% agarose gel (Pulsed Field Certified Agarose, Bio-Rad) at 14°C for 24h, 3V/cm, initial switch time 0.5 s, final switch time 7.0 s. For length calibration, we added $^{32}P$-labelled size markers (1kb DNA ladder, New England Biolabs Inc., Ipswich, MA, USA; DNA Molecular Weight Marker XV, Roche Diagnostics, Basel, Switzerland). Gels were dried (gel dryer, Bio-Rad, model 538) at room temperature and hybridized overnight at 37°C with a $^{32}P$-endlabelled oligonucleotide $(5'\text{-CCCTAA-}3')_4$ that binds to the telomeric single-strand overhang of non-denatured DNA. Subsequently, unbound oligonucleotides were removed by washing the gel for 30 min at 37°C with 0.25x saline-sodium citrate buffer. The radioactive signal of the sample-specific TL distribution was detected by a phosphor screen (MS, Perkin-Elmer Inc., Waltham, MA, USA), exposed overnight, and visualized using a phosphor imager (Cyclone Storage Phosphor System, Perkin-Elmer Inc.). We calculated average TL using ImageJ (v. 1.38x) as described by Salomons *et al.* (2009). For each sample the limit at the side of the short telomeres of the distribution was lane-specifically set at the point of the lowest signal (i.e. background intensity). The limit on the side of the long telomeres of the distribution was set lane-specifically where the signal dropped below Y, where Y is the sum of the background intensity plus 10 % of the difference between peak intensity and background intensity.

**Table S1.** Summary of the jackdaw pedigree pruned for telomere data information (R package PEDANTICS, Morrissey and Wilson 2010). Included are individuals with known early-life telomere length (TL) **(A)** at the age of 4 days (n=715) or **(B)** at the age of 29 days (n=474, a subset of individuals from A) and additionally individuals related to two or more of those phenotyped individuals.

| **Pedigree statistics** | **(A) Quantity (TL age 4 days)** | **(B) Quantity (TL age 29 days)** |
|---|---|---|
| Individuals | 1007 | 714 |
| Maternity links | 724 | 492 |
| Paternity links | 720 | 488 |
| pair-wise full sib relationships | 1289 | 787 |
| Maternal sibs | 1770 | 1022 |
| Paternal sibs | 1660 | 987 |
| Mothers | 174 | 133 |
| Fathers | 177 | 134 |
| Maternal grandmothers | 149 | 104 |
| Maternal grandfathers | 149 | 99 |
| Paternal grandmothers | 158 | 103 |
| Paternal grandfathers | 155 | 97 |
| Maximum pedigree depth | 6 | 6 |
| N pairwise relatedness: | | |
| >= 0.5 | 2733 | 1767 |
| >= 0.25 - < 0.5 | 2574 | 1438 |
| >= 0.125 - < 0.25 | 1773 | 966 |
| >= 0.025 - < 0.125 | 899 | 427 |

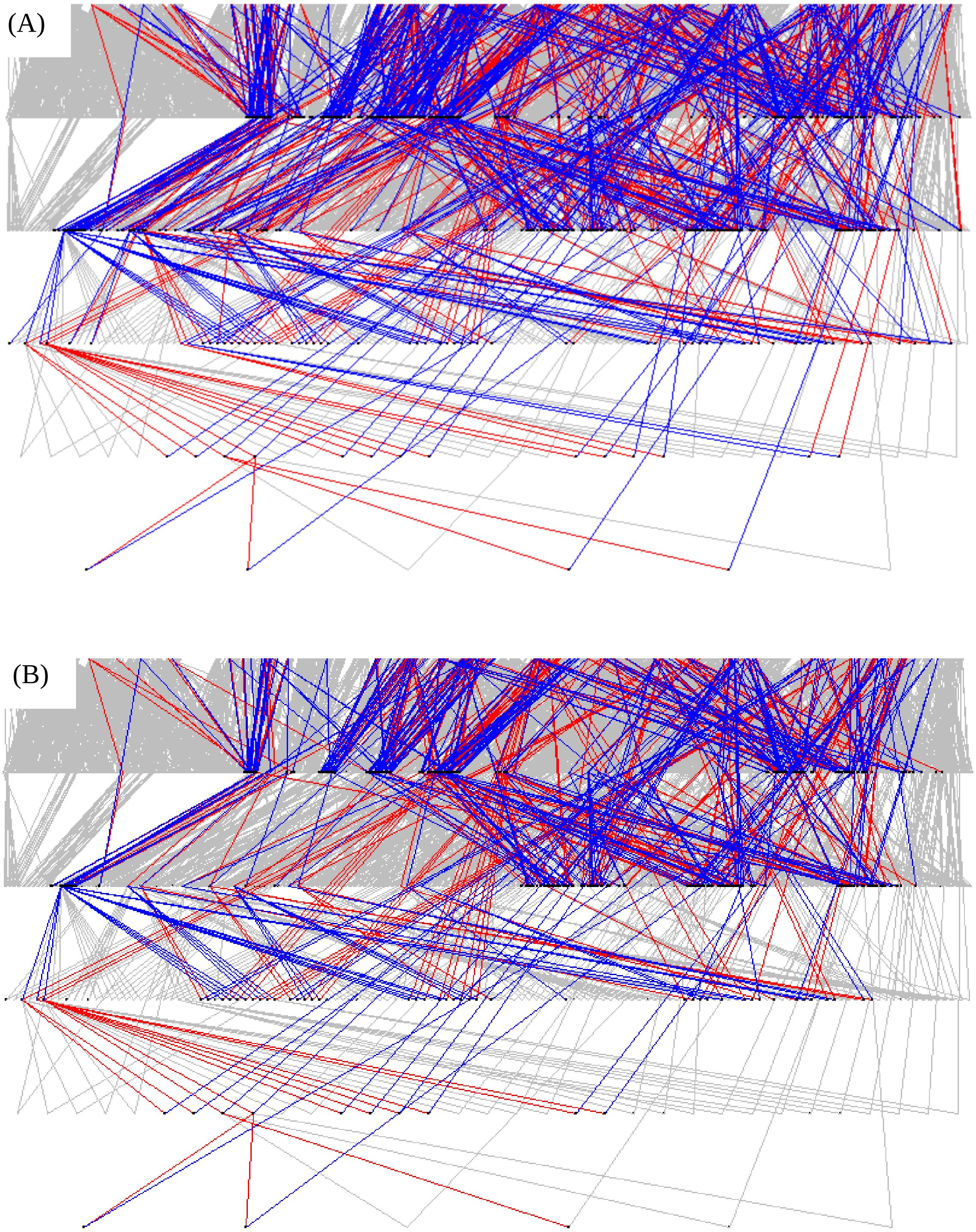

**Fig. S1. Pedigrees of the jackdaw population.** Pedigree **(A)** based on individuals phenotyped for telomere length at the age of 4 days and **(B)** at the age of 29 days, and each including respective relationship informative individuals. Every line connects a parent with one of its offspring. At the top of this six-generational pedigree are the founder individuals, at the bottom the 6$^{th}$ generation. Blue lines indicate informative paternal links, red lines indicate informative maternal links and overlay the full population pedigree, which is indicated in grey. For quantities and statistics on the pruned pedigrees see table S4. The pedigree images were generated using the R package PEDANTICS (Morrissey & Wilson 2010).

### *Animal model analyses*

**Table S2.** Univariate animal model with telomere length (in bp) at age 4 days as dependent variable, investigating maternal effects (mother ID) while controlling for father's age (n = 715 individuals).

| **Fixed effects** | **Coefficient (SE)** | | ***F*** | **df** | ***P* value** |
|---|---|---|---|---|---|
| Intercept | 7129.8 (77.3) | | | | |
| Father's age | -26.236 (14.040) | | 3.492 | 1, 251.2 | 0.063 |
| **Random effects** | **Variance (SE)** | **Proportion of variance (SE)** | $\chi^2$ | **df** | ***P* value** |
| Additive genetic | 182405.7 (43274.7) | 0.541 (0.116) | 16.317 | 1 | <0.001 |
| Mother ID | 14283.1 (19044.4) | 0.042 (0.056) | 0.531 | 1 | 0.466 |
| Brood ID (B1) | 15225.6 (14368.0) | 0.045 (0.043) | 1.387 | 1 | 0.239 |
| Birth year | 23470.3 (16762.7) | 0.070 (0.047) | 5.622 | 1 | 0.018 |
| Gel ID | 48499.2 (14118.8) | 0.144 (0.039) | 43.637 | 1 | <0.001 |
| Residual | 53203.9 (23127.4) | 0.158 (0.074) | | | |
| | | **Heritability $h^2$ (SE)** | | | |
| | | 0.632 (0.133) | | | |

Father ID as random effect could not be estimated and was therefore not included in the final model.

**Table S3.** Univariate animal model with telomere length (in bp) at age 4 days as dependent variable, with maternal effects (mother ID) while not controlling for father's age (n = 715 individuals).

| **Fixed effects** | **Coefficient (SE)** | | ***F*** | **df** | ***P* value** |
|---|---|---|---|---|---|
| Intercept | 7051.0 (64.2) | | - | - | - |
| **Random effects** | **Variance (SE)** | **Proportion of variance (SE)** | $\chi^2$ | **df** | ***P* value** |
| Additive genetic | 194500.4 (43440.6) | 0.573 (0.114) | 18.118 | 1 | <0.001 |
| Mother ID | 14037.5 (18569.1) | 0.041 (0.054) | 0.561 | 1 | 0.454 |
| Brood ID (B1) | 14606.5 (14129.8) | 0.043 (0.042) | 1.326 | 1 | 0.250 |
| Birth year | 22220.8 (16236.8) | 0.065 (0.045) | 4.889 | 1 | 0.027 |
| Gel ID | 47226.0 (13942.8) | 0.139 (0.039) | 41.416 | 1 | <0.001 |
| Residual | 47069.2 (23163.7) | 0.139 (0.073) | | | |
| | | **Heritability $h^2$ (SE)** | | | |
| | | 0.665 (0.130) | | | |

Father ID as random effect could not be estimated and was therefore not included in the final model.

**Table S4.** Univariate animal model with telomere length (in bp) at age 4 days as dependent variable, investigating paternal effects (father ID) while controlling for father's age (n = 715 individuals).

| **Fixed effects** | **Coefficient (SE)** | | ***F*** | **df** | ***P* value** |
|---|---|---|---|---|---|
| Intercept | 7128.7 (77.7) | | | | |
| Father's age | -26.148 (13.839) | | 3.57 | 1, 280.5 | 0.060 |
| **Random effects** | **Variance (SE)** | **Proportion of variance (SE)** | $\chi^2$ | **df** | ***P* value** |
| Additive genetic | 196091.3 (37481.8) | 0.582 (0.096) | 34.753 | 1 | <0.001 |
| Brood ID (B1) | 20619.0 (13313.2) | 0.061 (0.040) | 3.122 | 1 | 0.078 |
| Birth year | 24832.7 (17344.4) | 0.074 (0.049) | 6.045 | 1 | 0.014 |
| Gel ID | 47956.8 (13983.2) | 0.142 (0.039) | 43.319 | 1 | <0.001 |
| Residual | 47222.8 (20619.3) | 0.140 (0.066) | | | |
| | | **Heritability $h^2$ (SE)** | | | |
| | | 0.679 (0.109) | | | |

Father ID as random effect could not be estimated and was therefore not included in the final model.

**Table S5.** Univariate animal model with telomere length (in bp) at age of 29 days as dependent variable, controlling for father's age (n=474 individuals).

| **Fixed effects** | **Coefficient (SE)** | | ***F*** | **df** | ***P* value** |
|---|---|---|---|---|---|
| Intercept | 6868.3 (91.9) | | | | |
| Father's age | -33.5 (16.5) | | 4.1 | 1,285.1 | 0.043 |
| **Random effects** | **Variance (SE)** | **Proportion of variance** | $\chi^2$ | **df** | ***P* value** |
| Additive genetic | 279871.8 (54704.6) | 0.769 | 37.382 | 1 | <0.001 |
| Brood ID (B2) | 4989.7 (12600.4) | 0.014 | 0.084 | 1 | 0.681 |
| Birth year | 21031.8 (20946.8) | 0.058 | 1.268 | 1 | 0.111 |
| Gel ID | 45587.3 (16564.4) | 0.125 | 11.418 | 1 | <0.001 |
| Residual | 12273.3 (31227.1) | 0.034 | | | |
| | | **Heritability $h^2$ (SE)** | | | |
| | | 0.879 (0.125) | | | |

Mother ID and father ID as random effects could not be estimated and were therefore not included in the final model.

***Parent-offspring regressions***

**Table S6.** Father-offspring regression, telomere length measured at the age of 4 days in all (in bp), father's age not included (n=111 individual offspring of 28 fathers).

| **Fixed effects** | **Coefficient (SE)** | | ***F*** | **df** | ***P* value** |
|---|---|---|---|---|---|
| Intercept | 5967.2 (731.7) | | | | |
| Father's TL (age 4 days) | 0.181 (0.104) | | 3.009 | 1, 26 | 0.095 |
| **Random effects** | **Variance (SE)** | **Proportion of variance (SE)** | $\chi^2$ | **df** | ***P* value** |
| Father ID | 53544.1 (43258.3) | 0.160 (0.118) | 1.7458 | 1 | 0.186 |
| Brood ID (B1) | 32832.2 (47270.4) | 0.098 (0.143) | 0.4006 | 1 | 0.527 |
| Birth year | 41044.6 (42704.9) | 0.122 (0.115) | 2.1076 | 1 | 0.147 |
| Gel ID | 6945.5 (24253.4) | 0.021 (0.072) | 0.0844 | 1 | 0.771 |
| Residual | 200744.9 (41041.1) | 0.599 (0.136) | | | |

**Table S7.** Father-offspring regression, telomere length measured at the age of 4 days in offspring and at age at conception in fathers (n=352 individual offspring of 82 fathers).

| **Fixed effects** | **Coefficient (SE)** | | ***F*** | **df** | ***P* value** |
|---|---|---|---|---|---|
| Intercept | 5278.7 (423.0) | | | | |
| Father's TL (at conception) | 0.305 (0.073) | | 17.26 | 1, 95.7 | <0.001 |
| **Random effects** | **Variance (SE)** | **Proportion of variance (SE)** | $\chi^2$ | **df** | ***P* value** |
| Father ID | 70349.6 (27092.8) | 0.209 (0.073) | 8.596 | 1 | 0.003 |
| Brood ID (B1) | 28164.3 (22472.1) | 0.083 (0.067) | 2.272 | 1 | 0.132 |
| Birth year | 12578.8 (14778.7) | 0.037 (0.043) | 1.340 | 1 | 0.247 |
| Gel ID | 52655.3 (22383.3) | 0.156 (0.062) | 8.364 | 1 | 0.004 |
| Residual | 173611.3 (17608.1) | 0.515 (0.062) | | | |

**Table S8**. Parent-offspring regressions comparing telomere length change (Δ TL) between the ages of 4 and 29 days (in bp) of **(A)** mothers and their offspring (n=45 individual offspring of 14 mothers) and **(B)** fathers and their offspring (n=45 individual offspring of 12 fathers).

| **A** | | | | | |
|---|---|---|---|---|---|
| **Fixed effects** | **Coefficient (SE)** | | ***F*** | **df** | ***P* value** |
| Intercept | -168.8 (83.6) | | | | |
| Mother's ΔTL | 0.179 (0.235) | | 0.58 | 1, 7.55 | 0.469 |
| **Random effects** | **Variance** | **Proportion of variance** | $\chi^2$ | **df** | ***P* value** |
| Mother ID | 2883 | 0.079 | 0.334 | 1 | 0.563 |
| Birth year | 2533 | 0.069 | 0.280 | 1 | 0.597 |
| Gel ID | 13460 | 0.367 | 4.477 | 1 | 0.034 |
| Residual | 17772 | 0.485 | | | |
| **B** | | | | | |
| **Fixed effects** | **Coefficient (SE)** | | ***F*** | **df** | ***P* value** |
| Intercept | -253.9 (51.8) | | | | |
| Father's ΔTL | 0.038 (0.174) | | 0.05 | 1, 3.61 | 0.839 |
| **Random effects** | **Variance** | **Proportion of variance** | $\chi^2$ | **df** | ***P* value** |
| Father ID | 9187 | 0.328 | 0.888 | 1 | 0.346 |
| Birth year | 2738 | 0.098 | 1.206 | 1 | 0.272 |
| Gel ID | 6483 | 0.232 | 1.095 | 1 | 0.295 |
| Residual | 9571 | 0.342 | | | |

Brood ID (B2) as random effect could not be estimated and was therefore not included in the final models.

### References

Morrissey, M. B., & Wilson, A. J. (2010) PEDANTICS: an R package for pedigree-based genetic simulation and pedigree manipulation, characterization and viewing. *Molecular Ecology Resources*, *10,* 711-719. doi: 10.1111/j.1755-0998.2009.02817.x
